# Short communication: Antibiotic resistance in Gram-negative bacteria isolated from street-vended foods in Bangladesh

**DOI:** 10.1101/2022.06.28.497961

**Authors:** Fariha Chowdhury Meem, Topu Raihan, Jahid Hasan Shourove, Abul Kalam Azad, GM Rabiul Islam

**Author notes:** Corresponding Author: (GMRI).

## Abstract

Antibiotic-resistant pathogens disseminated through food are a public health concern. Although a significant proportion of the urban population in developing countries consume street-vended foods, their role in spreading antibiotic resistance has been rarely investigated. In this study, we evaluated the antibiotic resistance patterns of bacterial isolates (n = 50) collected from five categories of street-vended foods (phuchka, chatpati, sausage, bun, and salad) in Bangladesh. Antibiotic susceptibility to twelve antibiotics was investigated by the Kirby-Bauer disk diffusion method. We found a high prevalence of *E. coli* (n = 32) in street-vended foods, with most isolates (65.63%) exhibiting multidrug resistance. The multiple antibiotic resistance (MAR) index showed that 22 isolates had MAR above 0.2, with resistance mostly against oxacillin, ampicillin, and cefuroxime. From the rest, three representative isolates were selected for molecular identification by DNA sequencing of 16S rDNA. *Klebsiella oxytoca* showed multiple drug resistance (MDR) and was resistant to ampicillin, oxacillin, cefuroxime, and kanamycin. *Burkholderia fungorum* showed no distinct inhibition zone against ampicillin and chloramphenicol. Additionally, the *Serratia nematodiphila* isolate showed no distinct inhibition zone against three antibiotics, including ampicillin, oxacillin, and cefuroxime. These findings might contribute to the knowledge of emerging antibiotic-resistant foodborne pathogens and raise concerns about the safety of street-vended foods in Bangladesh.

## Introduction

Antibiotic-resistant (ABR) bacterial infections might become the leading cause of mortality worldwide by 2050 [1]. The resistant strains of foodborne pathogens significantly contribute to the scenario. The ABR bacteria are transmitted to humans through the food chain due to the uncontrolled use of antibiotics to promote growth in livestock, aquaculture, and apiculture [2, 3]. Among all the WHO regions, countries in Southeast Asia, including Bangladesh, have the highest risk of antibiotic resistance [4].

Street-vended food (SVF) is extremely popular among city dwellers as it is easily available, convenient, and inexpensive. SVF provides food security and employment to a significant proportion of the population in many developing countries. However, street food vendors generally lack basic hygienic practices and infrastructure, as well as have limited factual knowledge and poor understanding of food safety procedures [5]. Foodborne bacterial pathogens in SVF cause outbreaks of foodborne diseases like cholera, diarrhea, food poisoning, and typhoid fever [5]. In Bangladesh, foodborne diseases affect over 30 million individuals annually [6]. Furthermore, antibiotic-resistant bacteria present in various foods are a major cause of foodborne disease transmission, making the prevention of foodborne diseases very difficult. Although some studies in Bangladesh determined the antibiotic resistance levels in SVF, the information gathered in those studies was insufficient to understand the seriousness of the situation in Bangladesh [7-11]. Thus, more information is required to better understand the risk of exposure to ABR through food items, particularly SVFs. Thus, in this study, we assessed the antibiotic susceptibility of Gram-negative bacteria isolated from five different SVFs in Bangladesh.

## Materials and methods

### Sample collection and isolation of bacteria

In this study, 25 samples of five categories of SVFs (phuchka, chatpati, sausage, bun, and salad) were collected from street vendors. Approximately 200–250 g of food sample was collected in sterile bags, transported immediately to the laboratory, and stored at 4 °C until they were analyzed; samples were not stored for more than 12 h. All samples were inoculated on MacConkey agar plates and incubated at 37 °C for 24 h. Two suspected colonies of gram-negative bacteria with typical morphological characteristics and size were selected from each sample and isolated by the streak plate method. Furthermore, the Eosin Methylene Blue (EMB) agar medium was used to study the morphological characteristics of the suspected coliform bacteria, and the IMViC test was performed for biochemical identification. The experimental design of this study is shown in Fig 1.

**Fig 1.**
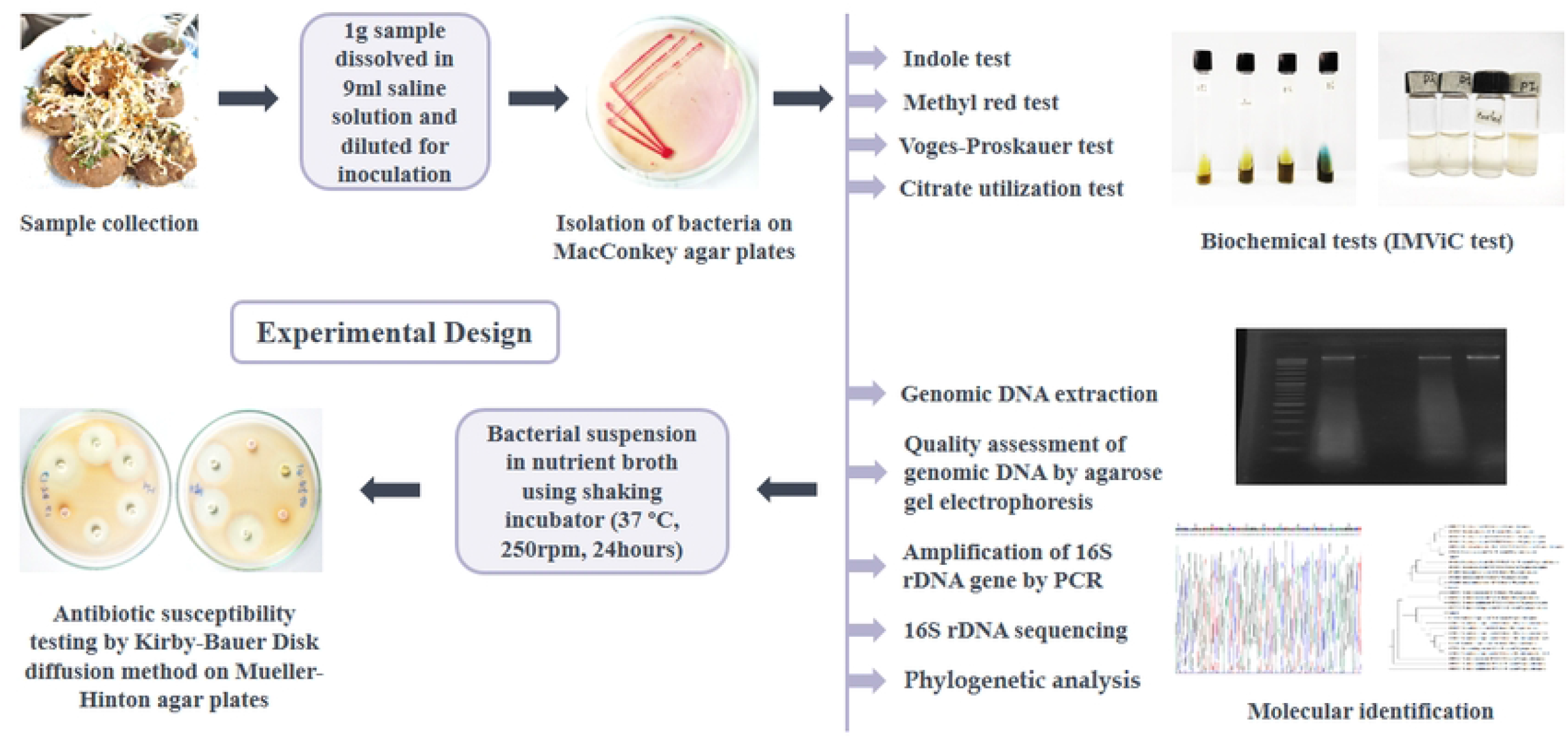
The experimental design.

### Molecular identification

Among the other isolates that could not be identified through biochemical tests, three distinct and representative isolates (F38, F42, and F45), with two or more 0 mm inhibition zone in the disk diffusion test, were selected for molecular identification by DNA sequencing of the 16S rDNA. Genomic DNA of the gram-negative bacteria was extracted using the Monarch DNA Gel Extraction Kit (New England Biolabs). After isolation of the genomic DNA, electrophoresis was performed in a horizontal gel apparatus (MyRun Cosmo Bio Co. Ltd., IMR-201). Electrophoresis was conducted in horizontal agarose gel (0.7%) to visually confirm the integrity of the genomic DNA [12]. PCR amplification of the bacterial 16S rDNA gene was performed using the forward primer 16S 8F (5′-AGA GTT TGA TCC TGG CTC AG-3′; 1500 bp) and the reverse primer 16S 1492R (5′-CGG TTA CCT TGT TAC GAC TT-3′; 1500 bp) [13]. In a thermal cycler (SimpliAmpTM Thermal Cycler, Applied Biosystem®, USA), PCR was performed using a 25 µL reaction mixture. The cycling parameters were 94 °C for 2 min, followed by 35 cycles at 94 °C for 30 s, 55 °C for 30 s, and 72 °C for 3 min, with a final extension at 72 °C for 10 min for each primer pair. Finally, 16S rDNA sequencing was performed using a DNA Sequencer (Model 3130, Applied Biosystem® Automated Genetic Analyzer) (Fig 2), and the sequences were submitted to National Centre for Biotechnology Information (NCBI), GenBank. The 16S rDNA sequences were aligned and paralleled with the sequences of the corresponding bacteria in NCBI, GenBank by BLAST (https://blast.ncbi.nlm.nih.gov/Blast.cgi) search. The sequences were aligned with the similar sequences using Clustal Omega (http://www.ebi.ac.uk/Tools/msa/clustalo/), and a phylogenetic tree was constructed using the MEGA 6.1 software [14].

**Fig 2.**
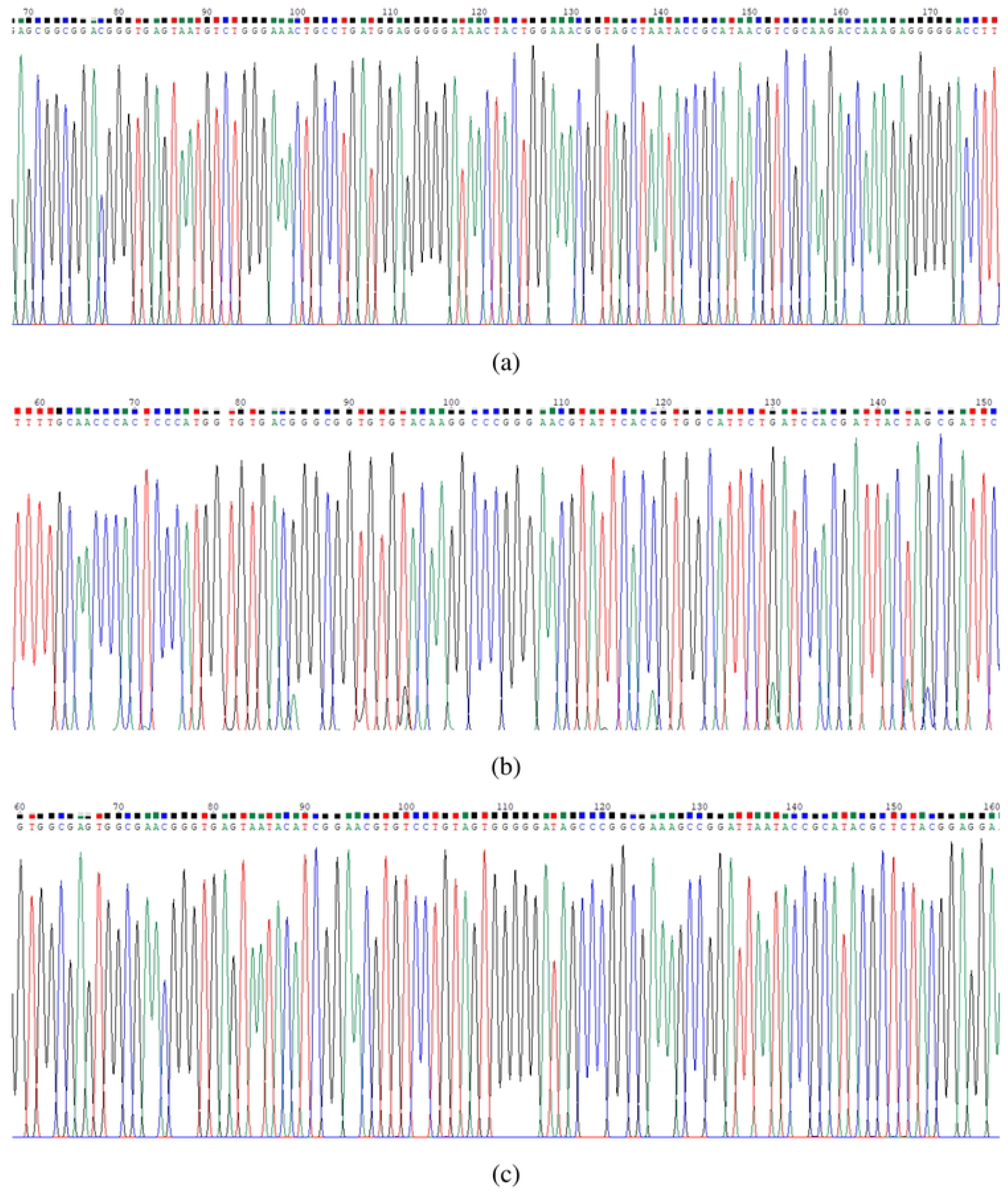
The chromatogram of the 16S rDNA gene sequence of (a) Isolate F38, (b) Isolate F42, and (c) Isolate F45 generated by the primers.

### Antimicrobial susceptibility testing

The Kirby-Bauer disk diffusion method was used to test antibiotic susceptibility after culturing the bacteria on Mueller Hinton agar (HiMedia; [7]). We applied 12 antibiotic agents (HiMedia), including Ampicillin (25 µg), Aztreonam (30 µg), Ceftriaxone (30 µg), Cefuroxime (30 µg), Chloramphenicol (30 µg), Cotrimoxazole or Trimethoprim/sulfamethoxazole (25 µg), Enrofloxacin (5 µg), Gentamicin (10 µg), Kanamycin (30 µg), Nalidixic Acid (30 µg), Oxacillin (1 µg), and Tetracycline (30 µg). Following the guidelines of the Clinical and Laboratory Standards Institute (CLSI), the inhibition zones were assessed and classified as sensitive or resistant [15]. Multidrug-resistant (MDR) isolates were defined as those that were resistant to at least one antimicrobial agent in three or more categories [16]. The MAR (multiple antibiotic resistance) index was calculated as the ratio of total antibiotics used to the number of antibiotics to which the bacterial isolate was resistant [17].

## Results

Among 50 colonies, 32 colonies (64%) showed green metallic sheen on EMB agar plates indicating *E. coli*, which was confirmed through biochemical tests. The phylogenetic tree constructed with similar sequences showed that the isolates, F38, F42, and F45, were closely related to the *Klebsiella oxytoca* strain PF 42 (Accession number: KY614353.1), the *Serratia nematodiphila* strain XM7 (MT023384), and the *Burkholderia fungorum* strain TN (KJ933410), respectively (see Fig 3).

**Fig 3.**
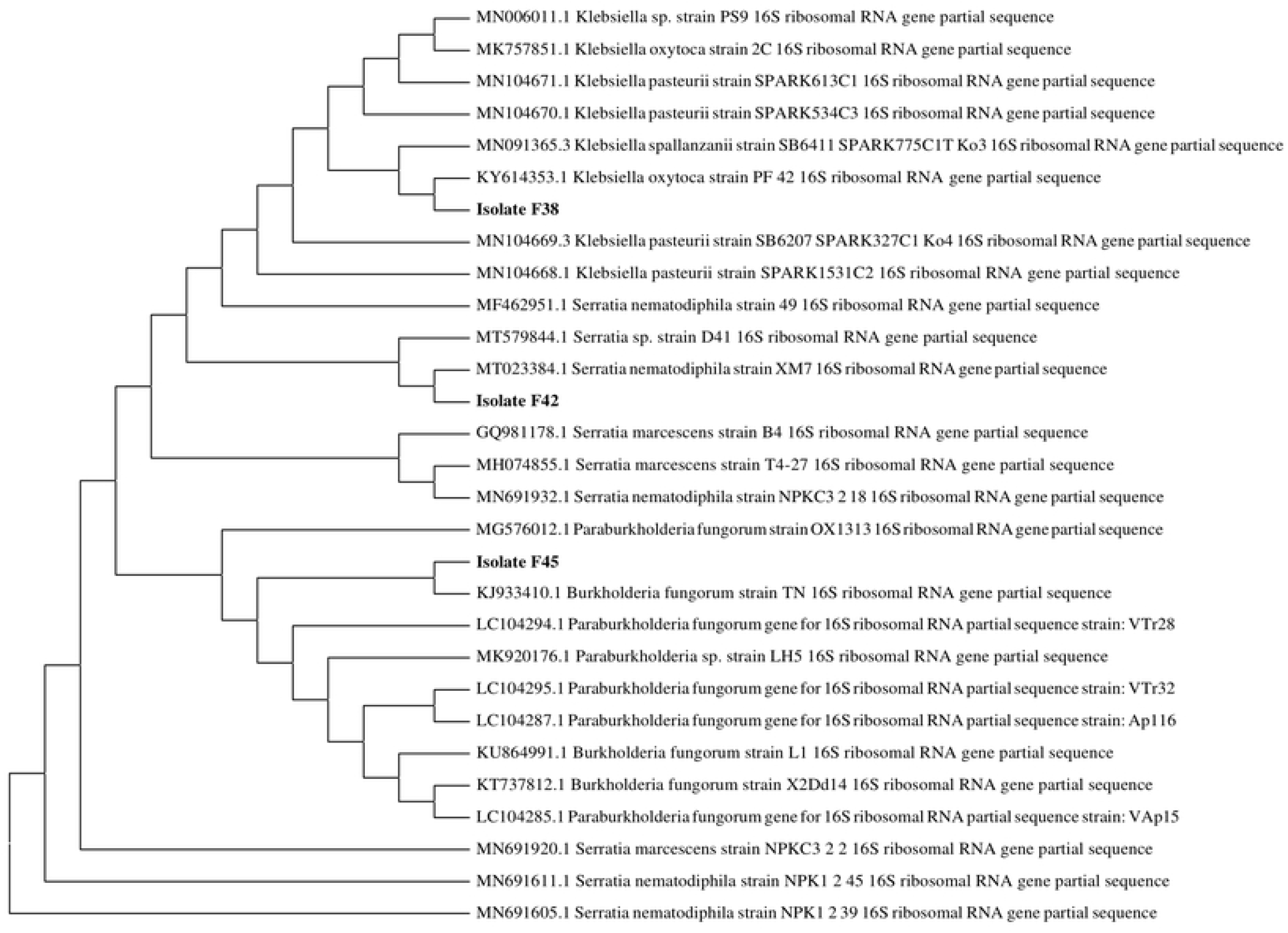
Phylogenetic tree.

The antibiotic susceptibility pattern showed that most of the *E. coli* isolates (65.63%) were multi-drug resistant (MDR), with high sensitivity to aztreonam (93.75%), enrofloxacin (93.75%), gentamicin (84.38%), and kanamycin (84.38%), whereas, high resistance was observed for the other antibiotics tested. Two isolates (F12 and F19) showed resistance to all the antibiotics (MAR index 1.0). Furthermore, among the MDR isolates, 100% were resistant to oxacillin, 95.23% were resistant to ampicillin, and 80.95% were resistant to cefuroxime. Twenty-two isolates (68.75%) had a MAR index greater than 0.2 and 10 (31.25%) isolates had a MAR index less than 0.2 (Table 1).

**Table 1:**
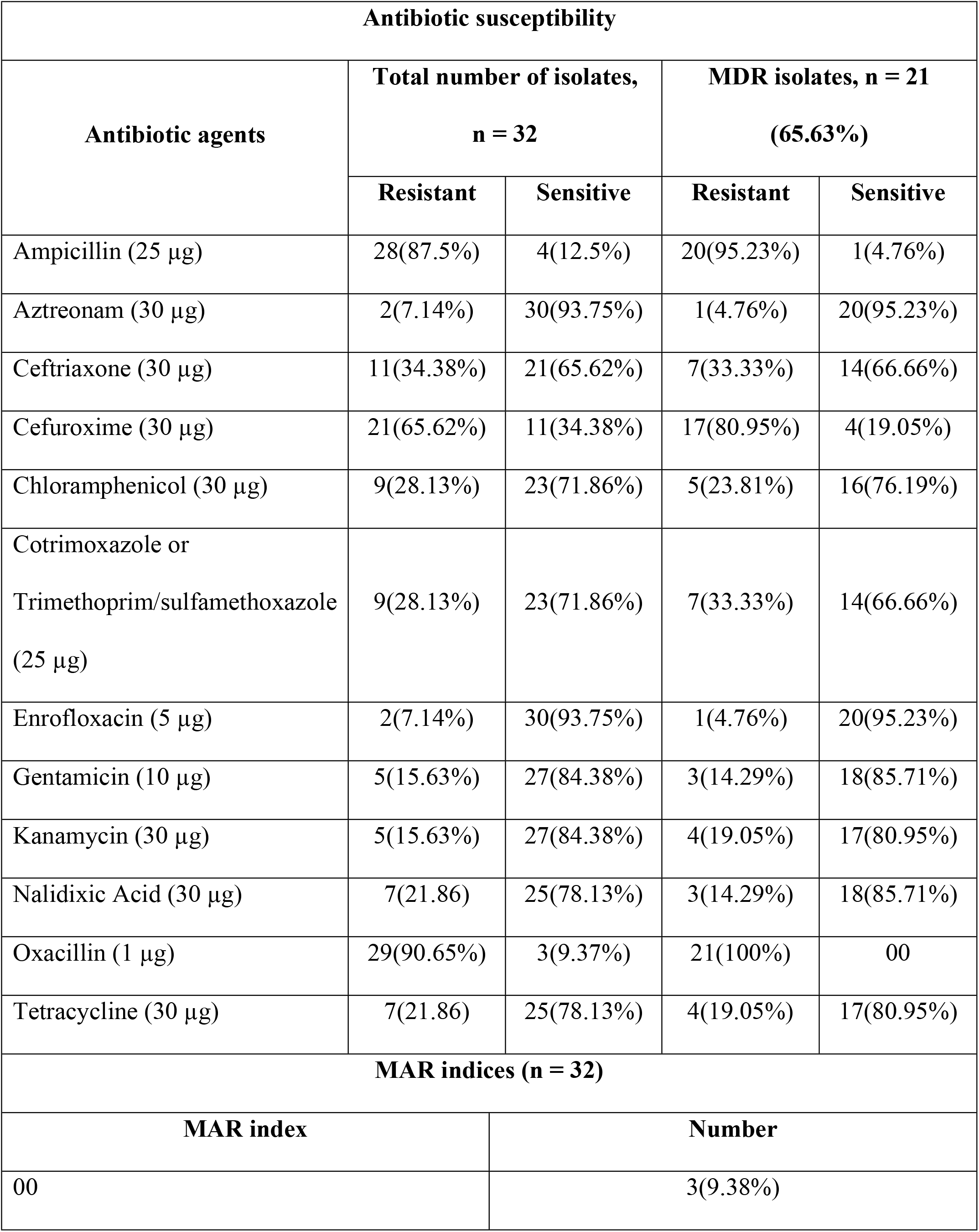
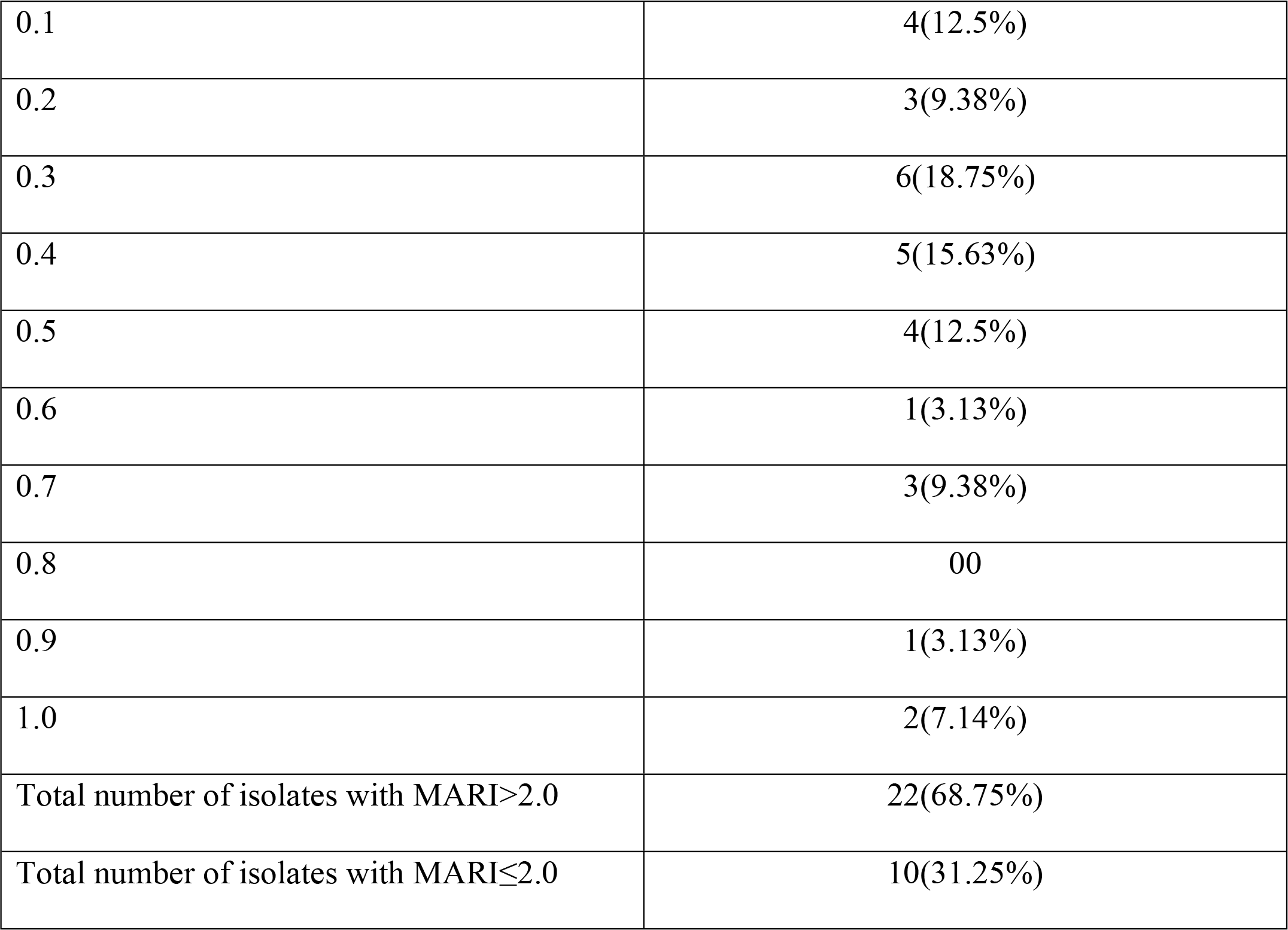
Antibiotic susceptibility pattern of *E. coli* isolates and the MAR index.

| **Antibiotic susceptibility** |  |  |  |  |
| --- | --- | --- | --- | --- |
| **Antibiotic agents** | **Total number of isolates,**<br>**n = 32** |  | **MDR isolates, n = 21**<br>**(65.63%)** |  |
|  | **Resistant** | **Sensitive** | **Resistant** | **Sensitive** |
| Ampicillin (25 µg) | 28(87.5%) | 4(12.5%) | 20(95.23%) | 1(4.76%) |
| Aztreonam (30 µg) | 2(7.14%) | 30(93.75%) | 1(4.76%) | 20(95.23%) |
| Ceftriaxone (30 µg) | 11(34.38%) | 21(65.62%) | 7(33.33%) | 14(66.66%) |
| Cefuroxime (30 µg) | 21(65.62%) | 11(34.38%) | 17(80.95%) | 4(19.05%) |
| Chloramphenicol (30 µg) | 9(28.13%) | 23(71.86%) | 5(23.81%) | 16(76.19%) |
| Cotrimoxazole or<br>Trimethoprim/sulfamethoxazole<br>(25 µg) | 9(28.13%) | 23(71.86%) | 7(33.33%) | 14(66.66%) |
| Enrofloxacin (5 µg) | 2(7.14%) | 30(93.75%) | 1(4.76%) | 20(95.23%) |
| Gentamicin (10 µg) | 5(15.63%) | 27(84.38%) | 3(14.29%) | 18(85.71%) |
| Kanamycin (30 µg) | 5(15.63%) | 27(84.38%) | 4(19.05%) | 17(80.95%) |
| Nalidixic Acid (30 µg) | 7(21.86) | 25(78.13%) | 3(14.29%) | 18(85.71%) |
| Oxacillin (1 µg) | 29(90.65%) | 3(9.37%) | 21(100%) | 00 |
| Tetracycline (30 µg) | 7(21.86) | 25(78.13%) | 4(19.05%) | 17(80.95%) |
| **MAR indices (n = 32)** |  |  |  |  |
| **MAR index** |  | **Number** |  |  |
| 00 |  | 3(9.38%) |  |  |

|  |  |
| --- | --- |
| 0.1 | 4(12.5%) |
| 0.2 | 3(9.38%) |
| 0.3 | 6(18.75%) |
| 0.4 | 5(15.63%) |
| 0.5 | 4(12.5%) |
| 0.6 | 1(3.13%) |
| 0.7 | 3(9.38%) |
| 0.8 | 00 |
| 0.9 | 1(3.13%) |
| 1.0 | 2(7.14%) |
| Total number of isolates with MARI>2.0 | 22(68.75%) |
| Total number of isolates with MARI≤2.0 | 10(31.25%) |

In our study, *Klebsiella oxytoca* isolate showed resistance against ampicillin, oxacillin, cefuroxime, and kanamycin. *Serratia nematodiphila* isolate did not show a clear inhibition zone against ampicillin, oxacillin, and cefuroxime. *Burkholderia fungorum* isolate did not show a clear inhibition zone against ampicillin and chloramphenicol (Table 2).

**Table 2:** The diameter of the disk diffusion inhibition zone.

| Antibiotic agents | Inhibition zone diameter (mm) |  |  |
| --- | --- | --- | --- |
|  | *Klebsiella oxytoca*<br>(Isolate F38) | *Serratia nematodiphila*<br>(Isolate F42) | *Burkholderia fungorum*<br>(Isolate F45) |
| Ampicillin (25 µg) | 00 (R) | 00 | 00 |
| Aztreonam (30 µg) | 21 (S) | 25 | 22 |
| Ceftriaxone (30 µg) | 22 (S) | 22 | 24 |
| Cefuroxime (30 µg) | 12 (R) | 00 | 11 |
| Chloramphenicol (30 µg) | 23 (S) | 28 | 00 |
| Cotrimoxazole or<br>Trimethoprim/sulfamethoxazole (25 µg) | 20 (S) | 21 | 28 |
| Enrofloxacin (5 µg) | 25 (S) | 28 | 27 |
| Gentamicin (10 µg) | 16 (S) | 20 | 11 |
| Kanamycin (30 µg) | 11 (R) | 11 | 20 |
| Nalidixic Acid (30 µg) | 22 (S) | 28 | 29 |
| Oxacillin (1 µg) | 00 (R) | 00 | 10 |
| Tetracycline (30 µg) | 16 (S) | 09 | 19 |

134 R =Resistant and S =Susceptible as per CLSI guidelines [15].

## Discussion

According to our knowledge, this is the first report on the prevalence of antibiotic-resistant *Klebsiella oxytoca* species in street food, as well as *Serratia nematodiphila*, and *Burkholderia fungorum* species in food items. All three isolates showed resistance against two or more antibiotics, including ampicillin. Similar results were found in a study on *Klebsiella oxytoca* isolates from animal-based food items and manured soil; which can be potential sources of contamination [18, 19]. In Bangladesh, the prevalence of *Klebsiella oxytoca* in street food has not been reported. However, a study on hospitalized patients in Bangladesh reported that *K. oxytoca* showed 100% resistance toward ampicillin, along with resistance to amoxicillin, ceftriaxone, ciprofloxacin, cotrimoxazole, and gentamicin by 75%, 25%, 50%, 50%, and 0%, respectively. The study found that gentamicin was the most potent drug [20]. The CLSI guidelines for determining antibiotic susceptibility using the disk diffusion method are currently unavailable for *Serratia nematodiphila* and *Burkholderia fungorum*. Very few studies have investigated the antibiotic resistance of *Serratia nematodiphila* and *Burkholderia fungorum*. A study on *Serratia* spp. isolated from sediments in Northeast India reported antibiotic resistance against five antibiotics [21]. Another study on isolates from the soil of the western Amazon reported that certain strains of *Burkholderia fungorum* exhibited resistance against at least four antibiotics [22]. Our study further highlighted the prevalence of multidrug-resistant *E. coli* in street foods of Bangladesh, considering that 64% of the isolates were identified as *E. coli*. The unhygienic food handling behaviors of street vendors might be the cause of fecal contamination, indicated by the presence of *E. coli* [23-26]. Additionally, 22 isolates had a MAR index >0.2, which indicated the overuse of antibiotics and the possibility of contamination from high-risk sources during production, handling of raw materials, manufacturing, and transportation [27-30].

Several studies conducted on street foods in Taiwan, the Philippines, Portugal, Mexico, and Ecuador reported resistance of *E. coli* isolates to ampicillin, chloramphenicol, tetracycline, sulfamethoxazole/trimethoprim, nalidixic acid, and gentamicin which support the results of our study [31-35]. To combat antibiotic resistance, the WHO has provided a list of priority pathogens, and members of Enterobacteriaceae (including *Klebsiella, E. coli, Serratia*, and *Proteus*) have been included in the most critical category [36]. Although our study had a limited number of isolates, the results suggested the prevalence and spread of lesser-known species of resistant bacteria throughout Bangladesh.

## Conclusion

Antibiotic-resistant microorganisms can spread through the food supply chain and adversely affect the ability to combat the persistent threat of infectious diseases. In this study, most of the *E. coli* isolates (90.63%) collected from street foods showed resistance against the applied antibiotics, which is alarming. Additionally, the presence of resistant strains of *Klebsiella oxytoca, Serratia nematodiphila*, and *Burkholderia fungorum* indicated potential health risks locally and globally. Future studies should cover a wider geographic area and focus on the horizontal transmission of multi-drug-resistant genes. Adequate measures, such as avoiding the use of unrestrained antibiotics and making street-vended foods more hygienic, should be taken to prevent the spread of resistant strains of pathogens. This study provides information regarding the emerging global issue of antibiotic-resistant bacteria. It emphasizes the need for developing standardized food safety measures and training street food vendors regularly.

## Acknowledgements

We would like to acknowledge all the staff of the Genetic Engineering and Biotechnology Department, Shahjalal University of Science and Technology, and Food Engineering and Tea Technology Department, Shahjalal University of Science and Technology.

The authors gratefully acknowledge the assistance of Biotech Concern with the DNA sequencing service.

## References

1. Cerqueira F, Matamoros V, Bayona JM, Berendonk TU, Elsinga G, Hornstra LM, et al. Antibiotic resistance gene distribution in agricultural fields and crops. A soil-to-food analysis. Environmental research. 2019;177:108608. doi: 10.1016/j.envres.2019.108608.

2. Caniça M, Manageiro V, Abriouel H, Moran-Gilad J, Franz CM. Antibiotic resistance in foodborne bacteria. Trends in Food Science & Technology. 2019;84:41–4. doi: 10.1016/j.tifs.2018.08.001.

3. Thapa SP, Shrestha S, Anal AK. Addressing the antibiotic resistance and improving the food safety in food supply chain (farm-to-fork) in Southeast Asia. Food Control. 2020;108:106809. doi: 10.1016/j.foodcont.2019.106809.

4. Hoque R, Ahmed SM, Naher N, Islam MA, Rousham EK, Islam BZ, et al. Tackling antimicrobial resistance in Bangladesh: A scoping review of policy and practice in human, animal and environment sectors. PLoS One. 2020;15(1):e0227947. doi: 10.1371/journal.pone.0227947.

5. Jahan M, Rahman M, Rahman M, Sikder T, Uson-Lopez RA, Selim ASM, et al. Microbiological safety of street-vended foods in Bangladesh. Journal of Consumer Protection and Food Safety. 2018;13(3):257–69. doi: 10.1007/s00003-018-1174-9.

6. Hasan M, Siddika F, Kallol MA, Sheikh N, Hossain MT, Alam MM, et al. Bacterial loads and antibiotic resistance profile of bacteria isolated from the most popular street food (Phuchka) in Bangladesh. Journal of Advanced Veterinary and Animal Research. 2021;8(3):361. doi: 10.5455/javar.2021.h523.

7. Mrityunjoy A, Kaniz F, Fahmida J, Shanzida J, Aftab UM, Rashed N. Prevalence of Vibrio cholerae in different food samples in the city of Dhaka, Bangladesh. International Food Research Journal. 2013;20(2).

8. Sarker N, Islam S, Hasan M, Kabir F, Uddin MA, Noor R. Use of multiplex PCR assay for detection of diarrheagenic Escherichia coli in street vended food items. Am J Life Sci. 2013;1(6):267–72. doi: 10.11648/j.ajls.20130106.15.

9. Tabashsum Z, Khalil I, Nazimuddin M, Mollah A, Inatsu Y, Bari ML. Prevalence of foodborne pathogens and spoilage microorganisms and their drug resistant status in different street foods of Dhaka city. Agriculture Food and Analytical Bacteriology. 2013;3(4):281–92.

10. Ahmed S, Tasnim UT, Pervin S, Islam M. An assessment of bacteriological quality of some fast food items available in Jessore city and antibiotic susceptibility of isolated Klebsiella spp. Int J Biosci. 2014;5:125–30. doi: 10.12692/ijb/5.9.125-8.

11. Ali M, Khan M, Saha ML. Antibiotic resistant patterns of bacterial isolates from ready-to-eat (RTE) street vended fresh vegetables and fruits in Dhaka City. Bangladesh Journal of Scientific Research. 2011;24(2):127–34.

12. Forcic D, Branovic-Cakanic K, Ivancic J, Jug R, Barut M, Strancar A. Purification of genomic DNA by short monolithic columns. Journal of Chromatography A. 2005;1065(1):115–20. doi: 10.1016/j.chroma.2004.10.030.

13. Iqbal A, Hakim A, Hossain MS, Rahman MR, Islam K, Azim MF, et al. Partial purification and characterization of serine protease produced through fermentation of organic municipal solid wastes by Serratia marcescens A3 and Pseudomonas putida A2. Journal of Genetic Engineering and Biotechnology. 2018;16(1):29–37. doi: 10.1016/j.jgeb.2017.10.011.

14. Wang X, Liu Y, Sun J. Physiological and Biochemical Characterization of Isolated Bacteria from a Coccolithophore Chrysotila dentata (Prymnesiophyceae) Culture. Diversity. 2021;14(1):2. doi: 10.3390/d14010002.

15. CLSI. Performance Standards for Antimicrobial Susceptibility Testing. Wayne, PA: Clinical and Laboratory Standards Institute: 2017.

16. Magiorakos A-P, Srinivasan A, Carey RB, Carmeli Y, Falagas M, Giske C, et al. Multidrug-resistant, extensively drug-resistant and pandrug-resistant bacteria: an international expert proposal for interim standard definitions for acquired resistance. Clinical microbiology and infection. 2012;18(3):268–81.

17. Titilawo Y, Sibanda T, Obi L, Okoh A. Multiple antibiotic resistance indexing of Escherichia coli to identify high-risk sources of faecal contamination of water. Environmental Science and Pollution Research. 2015;22(14):10969–80. doi: 10.1007/s11356-014-3887-3.

18. Abebe GM. Detection of Biofilm Formation and Antibiotic Resistance in Klebsiella Oxytoca and Klebsiella Pneumoniae from Animal Origin Foods. International Journal of Microbiology and Biotechnology. 2020;5(3):120.

19. Wang L, Zhao X, Wang J, Wang J, Zhu L, Ge W. Macrolide-and quinolone-resistant bacteria and resistance genes as indicators of antibiotic resistance gene contamination in farmland soil with manure application. Ecological Indicators. 2019;106:105456. doi: 10.1016/j.ecolind.2019.105456.

20. Chakraborty S, Mohsina K, Sarker PK, Alam MZ, Karim MIA, Sayem SA. Prevalence, antibiotic susceptibility profiles and ESBL production in Klebsiella pneumoniae and Klebsiella oxytoca among hospitalized patients. Periodicum biologorum. 2016;118(1). doi: 10.18054/pb.v118i1.3160.

21. Sarma B, Acharya C, Joshi S. Characterization of metal tolerant Serratia spp. isolates from sediments of uranium ore deposit of Domiasiat in Northeast India. Proceedings of the National Academy of Sciences, India Section B: Biological Sciences. 2016;86(2):253–60. doi: 10.1007/s40011-013-0236-0.

22. de Oliveira-Longatti SM, Marra LM, Soares BL, Bomfeti CA, Da Silva K, Ferreira PAA, et al. Bacteria isolated from soils of the western Amazon and from rehabilitated bauxite-mining areas have potential as plant growth promoters. World Journal of Microbiology and Biotechnology. 2014;30(4):1239–50. doi: 10.1007/s11274-013-1547-2.

23. Giri S, Kudva V, Shetty K, Shetty V. Prevalence and Characterization of Extended-Spectrum β-Lactamase-Producing Antibiotic-Resistant Escherichia coli and Klebsiella pneumoniae in Ready-to-Eat Street Foods. Antibiotics. 2021;10(7):850. doi: 10.3390/antibiotics10070850.

24. Rheinländer T, Olsen M, Bakang JA, Takyi H, Konradsen F, Samuelsen H. Keeping up appearances: perceptions of street food safety in urban Kumasi, Ghana. Journal of Urban Health. 2008;85(6):952–64.

25. Sarter G, Sarter S. Promoting a culture of food safety to improve hygiene in small restaurants in Madagascar. Food Control. 2012;25(1):165–71.

26. Rane S. Street vended food in developing world: hazard analyses. Indian journal of microbiology. 2011;51(1):100–6.

27. Ateba CN, Tabi NM, Fri J, Bissong MEA, Bezuidenhout CC. Occurrence of antibiotic-resistant bacteria and genes in two drinking water treatment and distribution systems in the North-West Province of South Africa. Antibiotics. 2020;9(11):745.

28. Ahmed HA, El Bayomi RM, Hussein MA, Khedr MH, Remela EMA, El-Ashram AM. Molecular characterization, antibiotic resistance pattern and biofilm formation of Vibrio parahaemolyticus and V. cholerae isolated from crustaceans and humans. International Journal of Food Microbiology. 2018;274:31–7.

29. Saeed E, Amer AAE-M, Keshta HG, Hafez EE, Sultan RM, Khalifa E. Prevalence, antibiotic sensitivity profile, and phylogenetic analysis of Escherichia coli isolated from raw dromedary camel milk in Matrouh Governorate, Egypt. Journal of Advanced Veterinary and Animal Research. 2022;9(1):138.

30. Maloo A, Fulke AB, Mulani N, Sukumaran S, Ram A. Pathogenic multiple antimicrobial resistant Escherichia coli serotypes in recreational waters of Mumbai, India: a potential public health risk. Environmental Science and Pollution Research. 2017;24(12):11504–17.

31. Lin L, Wang S-F, Yang T-Y, Hung W-C, Chan M-Y, Tseng S-P. Antimicrobial resistance and genetic diversity in ceftazidime non-susceptible bacterial pathogens from ready-to-eat street foods in three Taiwanese cities. Scientific reports. 2017;7(1):1–9. doi: 10.1038/s41598-017-15627-8.

32. Manguiat LS, Fang TJ. Microbiological quality of chicken-and pork-based street-vended foods from Taichung, Taiwan, and Laguna, Philippines. Food microbiology. 2013;36(1):57–62. doi: 10.1016/j.fm.2013.04.005.

33. Campos J, Gil J, Mourao J, Peixe L, Antunes P. Ready-to-eat street-vended food as a potential vehicle of bacterial pathogens and antimicrobial resistance: an exploratory study in Porto region, Portugal. International journal of food microbiology. 2015;206:1–6. doi: 10.1016/j.ijfoodmicro.2015.04.016.

34. Zurita J, Yánez F, Sevillano G, Ortega-Paredes D, Paz y Miño A. Ready-to-eat street food: a potential source for dissemination of multidrug-resistant Escherichia coli epidemic clones in Quito, Ecuador. Letters in applied microbiology. 2020;70(3):203–9. doi: 10.1111/lam.13263.

35. Estrada-Garcia T, Lopez-Saucedo C, Zamarripa-Ayala B, Thompson M, Gutierrez-Cogco L, Mancera-Martinez A, et al. Prevalence of Escherichia coli and Salmonella spp. in street-vended food of open markets (tianguis) and general hygienic and trading practices in Mexico City. Epidemiology & Infection. 2004;132(6):1181–4. doi: 10.1017/S0950268804003036.

36. Organization WH. WHO publishes list of bacteria for which new antibiotics are urgently needed. 2019. Available from: http://www.who.int/mediacentre/news/releases/2017/bacteria-antibiotics-needed/en.

